# Mutant strains of *Escherichia coli* lacking global regulators, *arcA* and *fis*, demonstrate better growth fitness by pathway reprogramming under acetate metabolism

**DOI:** 10.1101/2021.06.24.449759

**Authors:** Shikha Jindal, Mahesh S. Iyer, Poonam Jyoti, Shyam Kumar Masakapalli, K.V. Venkatesh

## Abstract

Global regulatory transcription factors play a significant role in controlling microbial metabolism under genetic and environmental perturbations. A systems-level effect of carbon sources such as acetate on microbial metabolism under disrupted global regulators has not been well established. Acetate is one of the substrates available in a range of nutrient niches such as the mammalian gut and high-fat diet. Therefore, investigating the study on acetate metabolism is highly significant. It is well known that the global regulators arcA and fis regulate acetate uptake genes in *E. coli* under glucose condition. In this study, we deciphered the growth and flux distribution of *E.coli* transcription regulatory knockout mutants Δ*arcA*, Δ*fis* and double deletion mutant, Δ*arcAfis* under acetate using ^13^C-Metabolic Flux Analysis which has not been investigated before. We observed that the mutants exhibited an expeditious growth rate (~1.2-1.6 fold) with a proportionate increase in acetate uptake rates compared to the wild-type. ^13^C-MFA displayed the distinct metabolic reprogramming of intracellular fluxes, which conferred an advantage of faster growth with better carbon usage in all the mutants. Under acetate metabolism, the mutants exhibited higher fluxes in the TCA cycle (~18-90%) and lower gluconeogenesis flux (~15-35%) with the proportional increase in growth rate. This study reveals a novel insight by stating the sub-optimality of the wild-type strain grown under acetate substrate aerobically. These mutant strains efficiently oxidize acetate to acetyl-CoA and therefore are potential candidates that can serve as a precursor for the biosynthesis of isoprenoids, biofuels, vitamins and various pharmaceutical products.

**Importance:** Unravelling the role of global regulatory genes on microbial metabolism of substrates available in various growth niche is important. Studies have shown that the global transcriptional regulators *arcA* and *fis*, under glucose availability, suppress the acetate uptake genes indicating a link between nutrient source and gene regulatory control. This work is focused on deciphering the influence of these regulators on acetate metabolism in *E.coli*. Growth studies using knockout strains (Δ*arcA,* Δ*fis and* Δ*arcAfis*) and ^13^C Metabolic flux analysis defined precise metabolic phenotypes under acetate metabolism. Interestingly, the mutants showed metabolic readjustment to facilitate optimal biomass requirements and a better balance between energy and precursor synthesis, resulting in better growth, which lacked in the wild-type strain. The outcomes of this study will be leveraged in understanding the regulatory control under various nutrient shifts.

## Introduction

Bacteria encounters diverse ranges of nutrient niches, facilitating the activation of regulatory machinery to maintain homeostasis (Fong and Palsson, 2004; Gfp, 2007; Tenaillon et al., 2012). Different metabolic pathways are prevalent for yielding energy in the organism to combat the nutrient stress under its preferred substrate limitation (Tramontano et al., 2018). In *E.coli*, glucose is a native carbon source, but the cell often confronts different substrates present in the environment (Rowland et al., 2018), such as acetate in the human gut systems, and ketogenic diet (Shiloach J et al., 2009). Introducing genetic perturbations and the altered carbon substrate into the cell allows one to understand the mechanism involved in the robustness of the biological systems. At the genetic level, global transcriptional regulators control more than half of all the genes that are extensively studied under glucose condition, but lesser investigated when acetate is employed as a carbon source. Among the global regulators, *fis and arcA* act as a repressor of the acetate uptake gene (Browning et al., 2004; Seshasayee et al., 2012; Iyer et al., 2020) and regulates various pathways such as the tricarboxylic acid (TCA cycle) and glyoxylate shunt under glucose metabolism aerobically (Iuchi and Lin, 1988; Perrenoud and Sauer, 2005; Park et al., 2013).

Acetate gets actively oxidized to acetyl-CoA by acetate synthetase (*acs*) under the aerobic condition at the expense of ATP, as shown in Fig. 1 (Kumari et al., 1995; Kumari et al., 2000; Wolfe, 2005). Furthermore, it enters the TCA cycle, glyoxylate shunt (El-Mansi et al., 2006) and to upper metabolic pathways such as gluconeogenesis (Christopher P. Long et al., 2017) and pentose phosphate (PP) pathway for its biomass and energy demands (Dolan and Welch, 2018). Previous studies have reported that under acetate metabolism, the gene expression of the TCA cycle genes, the glyoxylate shunt genes and gluconeogenesis genes are enhanced compared to the glucose condition (Shimizu, 2003). Potentially, intracellular fluxes can provide novel insights into the system associated with altered cellular demands of NADPH, ATP, and other cofactors by rerouting the fluxes, conferring its pleiotropic phenotypes (Zhao and Shimizu, 2003). Here, for the first time, we have elucidated the effect of the knockout of transcriptional regulators (*arcA* and *fis*) under acetate aerobic metabolism using a high-resolution ^13^C-Metabolic Flux Analysis (MFA) approach.

**Fig. 1.**
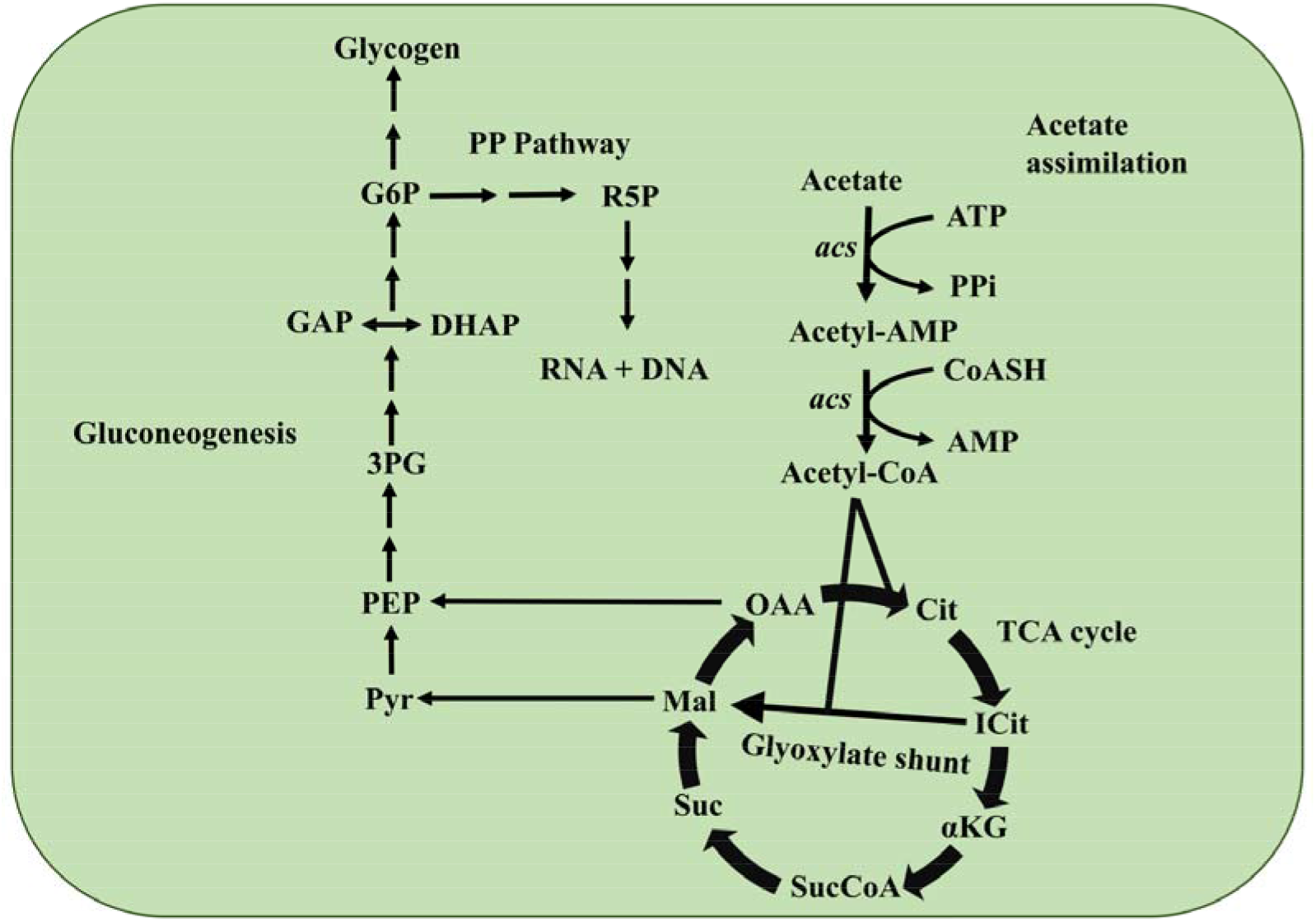
The metabolic network of the central carbon pathway describes the acetate assimilation in *E.coli* k-12 with the first step of acetate oxidation is catalyzed by acetyl synthetase (*acs*) followed by TCA cycle along with glyoxylate shunt. Further, the onset of the gluconeogenesis pathway begins from the PEP. From G6P, the flux bifurcates to the Pentose phosphate (PP) pathway. Continuous arrows denote more than one reaction in the pathway.

Interestingly, ^13^C-MFA sheds light on the metabolic reprogramming of fluxes in the knockout strains that gave an advantage towards their better growth compared to the wild-type strain. This revealed the novel insight of conferring optimality in mutants under acetate metabolism compared to the wild-type.

## Material and Methods

### Chemicals

All chemicals and M9 minimal media components used in this study were purchased from Sigma-Aldrich. ^13^C Acetate tracers were purchased from Cambridge Isotope laboratory: [1,2-^13^C] Acetate (99.5 atom% ^13^C) and [1-^13^C] Acetate (99.2% ^13^C).

### Bacterial strains and growth conditions

In this study, *E.coli* K-12 MG1655 (CGSC#6300) was used as the wild-type (WT) strain along with its mutant strains, Δ*fis*, Δ*arcA* and Δ*arcAfis,* which were constructed using a one-step inactivation protocol as mentioned in Supplementary material 1 (Datsenko and Wanner, 2000). These knockout strains were validated by PCR. WT and its mutant strains were cultivated aerobically in sodium acetate (4 g/L) as a sole carbon source in an M9 minimal media (Composition per litre of distilled water: 6 g Na2HPO4.12H2O, 3 g KH2PO4, 1 g NH4Cl, 0.5 g NaCl along with 1 g MgSO4 and 0.5 g CaCl2). A working volume of 5ml was used in each batch culture at 37°C on a rotary shaker (Eppendorf). The pH of the growth medium was set to 7 ± 0.05. All solutions were filter sterilized.

### ^13^C Tracer Experiment

Parallel labelling experiment was performed for each strain by using 4 g/L (3.54 mM) of ^13^C acetate (60% [1-^13^C] acetate + 40% [1, 2-^13^C] acetate). Cultures were harvested in the exponential phase. Each labelling experiment was performed in triplicates (Fig. 2).

**Fig. 2.**
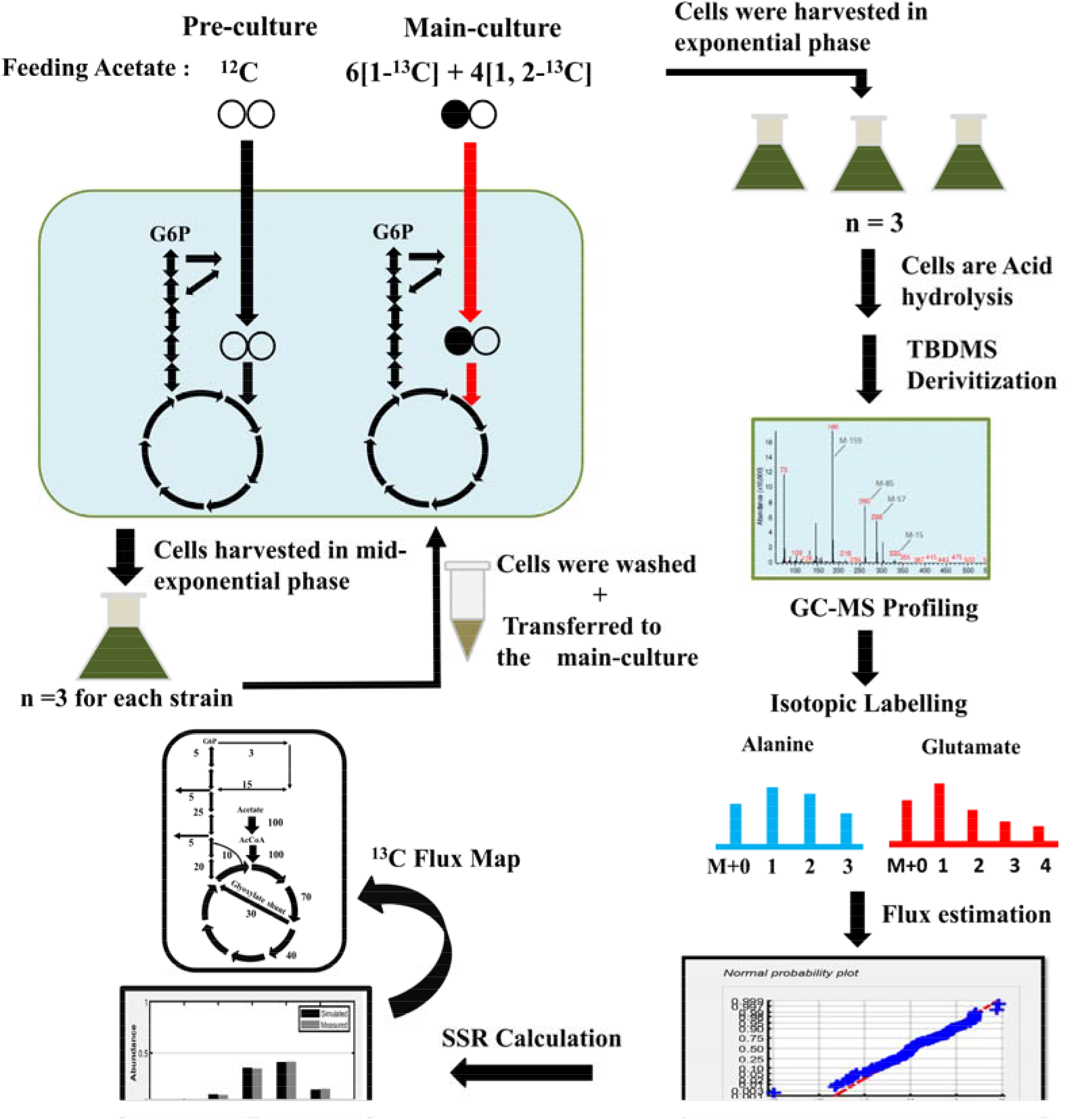
A schematic illustration of the workflow for the ^13^C-tracer experiment and its analysis are shown. Firstly, pre-culture was fed with unlabelled acetate and grown till mid-exponential phase. Thereafter, cells were harvested, washed and transferred into main-culture with 60%[1-^13^C] Acetate + 40% [1,2-^13^C] Acetate for all the strains in triplicates (n=3). These cells were harvested in exponential phase for hydrolysis and derivatization (TBDMS) to get proteinogenic amino-acids. Further, GC-MS profile was generated. With computational analysis, mass isotopomer distribution (MIDs) were baseline and mass corrected. These MIDs were incorporated into the INCA software for flux estimation. The flux map was produces with an acceptable range of SSRs.

### Measurement of Microbial Growth, biomass and CHNS analysis

Growth of *E.coli* WT, Δ*fis*, Δ*arcA* and Δ*arcAfis* cultures was monitored spectrophotometrically by measuring the optical density at 600 nm (OD_600_). For each strain, the growth rate was estimated based on linear regression of the natural logarithm of OD_600_ in the exponential phase versus time. Furthermore, the growth rate was utilized to calculate the acetate uptake rate and biomass yield (Chen et al., 2011). Cells were harvested at regular intervals in the exponential phase and lyophilized for the dry cell weight (DCW) and biomass quantification. To determine the relationship between OD_600_ and dry cell weight, OD_600_ was measured at regular intervals in the log phase with its corresponding dry cell weight per litre. Therefore, a conversion coefficient of 0.43 gDCW/ L / OD_600_ was calculated. To perform the elemental analysis of carbon and nitrogen, 10 mg of lyophilized cell pellet was subjected to CHNS (Carbon Hydrogen Nitrogen Sulphur) analyzer (ThermoFinnigan).

### 1H-NMR analysis of acetate

Acetate concentration was determined in the exponential growth phase by using ^1^H proton NMR (JNM-ECX500 spectrometer). The cell culture was centrifuged to remove the cell debris, and the supernatant was collected by filter sterilization. 300 μL of filter-sterilized supernatant was diluted with the 200 μL of double-distilled water (D_2_O). Further, 100 μL of 0.1% DSS (4,4-dimethyl-4-silapentane-1-sulfonic acid) was added to each sample as an internal standard. This 600ml solution was gently mixed and transferred to 5mm NMR glass tubes for acquiring acetate quantification using JOEI-DELTA. Each Spectra of ^1^H-NMR was recorded with 64 scans and a pulse width of 11.6 μs. The spectral intensities were calculated using the Delta software (version 5.7). The concentration of acetate was calculated at equal intervals in the exponential phase to calculate its consumption rate.

### Gas Chromatography-Mass Spectroscopy (GC-MS)

GC-MS measurements were executed on an Agilent 7890A GC system equipped with a DB-5ms capillary column (30 m, 0.25 mm, 0.25 μm-phase thickness; Agilent J&W Scientific), connected to a Waters Quattro Micro Tandem Mass Spectrometer (GC-MS/MS) operating under ionization by electron impact (EI) at 70 eV. An injection volume of 1 μL was used for all the samples to be analyzed. Helium, as a carrier gas, was used at 0.6 ml/min. The initial temperature of the oven was at 120°C held for 5 min. It was followed by 4°C/ min gradient upto 270°C, held for 3 min, followed by 20°C/min ramped to 320°C. The GC-MS spectra were recorded for a total run time of 50 min with a scanning range of 40-600 m/z.

### Sample preparation and derivatization for GC-MS

Bacterial cells (2 mg dry weight) were subjected to acid hydrolysis by adding 500 μl of 6 M HCl and incubating at 95°C for 16 h (Shree and Masakapalli, 2018). 20 μl of acid hydrolysate was subjected to a vacuum concentrator (Thermo-315 scientific, Waltham, MA, USA) for 3 h at 35°C to obtain the dried extract. To detect the amino acids, the dried extracts were subjected to TBDMS (tert-Butyldimethylsilyl chloride) derivatization. 30 μl of pyridine was added to each dried hydrolysate extract sample and incubated at 37°C for 30 min at 900 rpm on a thermomixer. Thereafter, 50 μl of MtBSTFA ((N-Methyl-N-(t-butyldimethylsilyl trifluoroacetamide)) + 1% t-BDMCS (N-methyl-N-(t-butyldimethylsilyl) trifluoroacetimide) + 1% t-butyl-dimethylchlorosilane was added to each sample and incubated at 60°C for 30 min at 900 rpm (thermomixer). Thereafter, samples were centrifuged at 13,000 × g for 12 min, and 50 μl of supernatant was transferred to GC-MS vials for injection.

### GC-MS Data analysis with statistics

The raw GC-MS data files were generated and the baseline corrections were performed using MetAlign Software (Lommen and Kools, 2012). Agilent ChemStation software was employed to obtain the intensity of the mass ions corresponding to each amino acid fragment (Masakapalli et al., 2014). Each derivatized amino acid was confirmed using the National Institute of Standards and Technology (NIST, Maryland) library and its corresponding standards. Thereafter, derivatised amino acid fragments were mass corrected for the natural ^13^C isotope abundance using IsoCorr software (Millard et al., 2012). The corrected mass isotopomer distribution (MID) of each stable amino acid fragment was selected (Supplementary materials 2) and the average ^13^C abundance calculated (Antoniewicz et al., 2007). For ^13^C-Metabolic flux distribution, the fitness of good (Antoniewicz et al., 2006) was determined by the acceptable range of the sum of square residues (SSRs).

### Metabolic network model and 13 C-MFA

The stoichiometry model of *E.coli* for 13C- FA was adapted from (Zhao and Shimizu, 2003) with a few modifications, such as the addition of reactions and cofactor balance, as mentioned in Supplementary materials 3. Modelling was performed using isotopomer network compartmental analysis (INCA) software (Young, 2014) which is based on the elementary metabolite units framework (Antoniewicz et al., 2007) in Matlab (MathWorks Inc., Natick, MA, USA). This metabolic network included all the major metabolic pathways such as the TCA cycle, the anaplerotic pathway, the glyoxylate shunt, the gluconeogenic pathway and the PP pathway, Biosynthetic reactions of amino acids, an oxidative phosphorylation reaction, a transhydrogenase reaction and a biomass formation reaction. New reactions along with cofactors which are a part of acetate metabolism, were included in the carbon transition network (Zhao and Shimizu, 2003), such as deamination of serine to pyruvate (Long et al., 2017a), atmospheric interference in CO2 dilution (Leighty and Antoniewicz, 2012), the reaction from G6P to the glycogen synthesis (Chen et al., 2010). Metabolic fluxes were assessed by minimizing the residual SSRs between the simulated and the experimental measured MIDs of the amino acids using non-linear least-squares regression (Long et al., 2017b). The accepted range of SSRs was chosen for each strain. Parameter optimization was performed to ensure that the best fit and flux boundaries were achieved.

### Statistical analysis

In each experiment, all data values are mentioned as mean ± Standard deviation (SD). Student’s t-test was performed to determine the statistical significance across WT, Δ*fis*, Δ*arcA* and Δ*arcAfis* in triplicates.

## Results

### Mutants exhibited a faster growth phenotype under acetate metabolism

Knockout strains, Δ*fis*, Δ*arcA* and Δ*arcAfis*, were constructed on *E.coli* MG1655 using homologous recombination. We monitored the physiological characteristics of each mutant relative to the WT in the exponential phase of growth with acetate as a carbon source (Fig. 3). All the mutants showed a higher growth rate (h^−1^) compared to WT (Fig. 3A). Δ*fis* had the highest growth rate of 0.26 h^−1^ which is about 50% higher than the WT with 0.16 h^−1^ (P < 0.01). On the other hand, Δ*arcA* (0.2 h^−1^) and Δ*arcAfis* (0.24 h^−1^) showed a 20% and 50% higher growth rate than the WT, respectively. Interestingly, it was also observed that the double mutant, Δ*arcAfis* showed a growth rate similar to that of Δ*fis*. The acetate uptake rate (Fig. 3B) demonstrated an upward trend similar to the growth rate in mutants, wherein Δ*fis* had the highest uptake rate (50%) followed by Δ*arcAfis* (47%) and Δ*arcA* (18%). However, the biomass yield of the mutants was statistically increased by 12% in Δ*fis*, 10% in Δ*arcAfis* and 4% in Δ*arcA* compared to the WT (Fig. 3C). Despite an increase in growth rate, and the acetate uptake rate in all the mutants, the ammonia uptake rate (Fig. 3D) was lower (~16-38%) compared to WT. Among the mutants, Δ*arcA* showed the lowest ammonia uptake (1.1 mmol/gDCW/h).

**Fig. 3.**
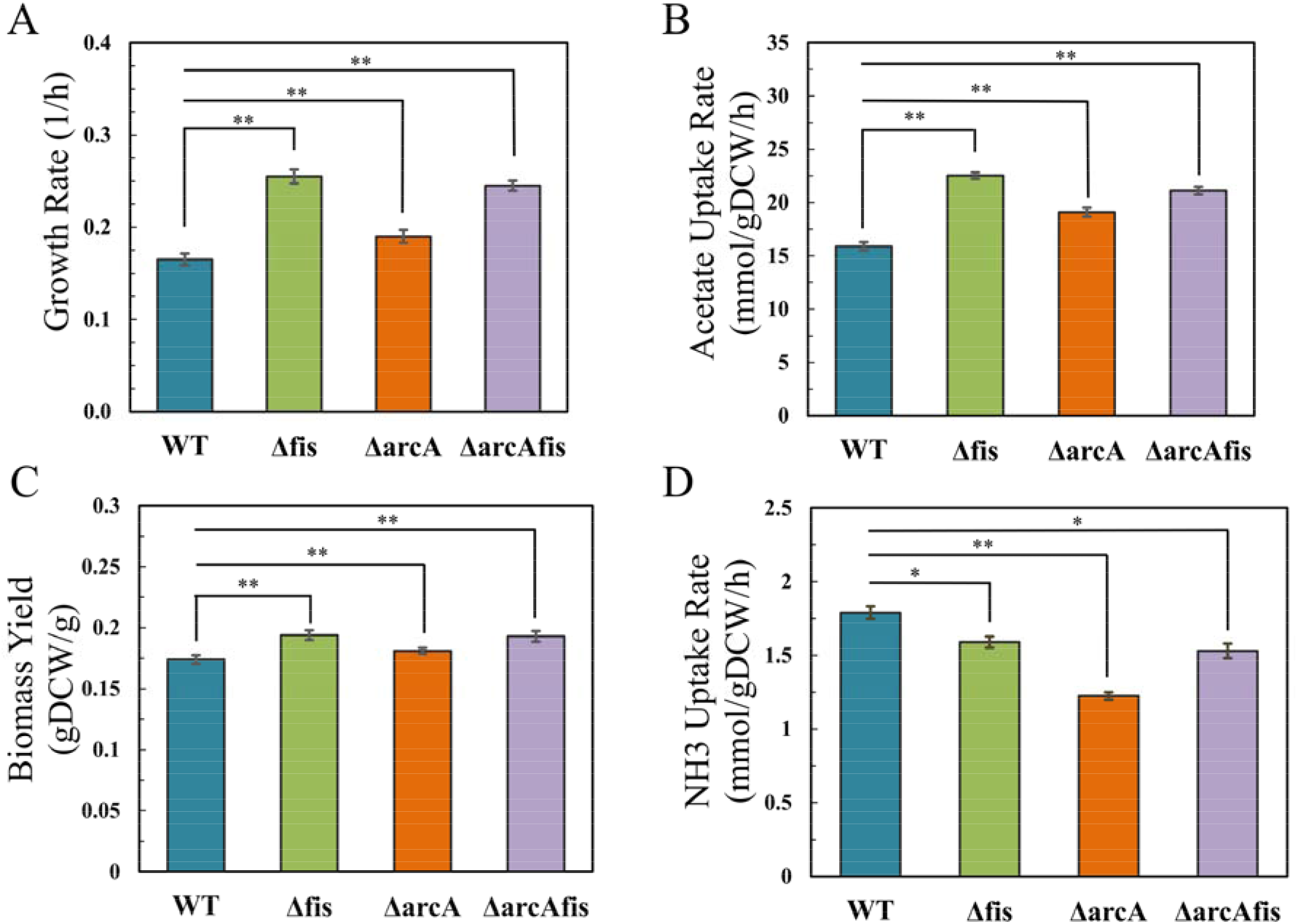
Physiological characteristics of *E.coli* WT (blue) and mutants Δ*fis* (green), Δ*arcA* (red) and Δ*arcAfis* (purple) grown under acetate minimal media A) Growth rates were measured in exponential phase B) Acetate uptake rate measured by ^1^H-NMR C) Total Biomass yields and D) Ammonia uptake rate in all the strains. Each experiment is performed in triplicates (n=3). Error bars represent the standard deviation derived from three replicates of each strain. Statistical significance is shown by asterisks. One asterisk (*) = P<0.05 and two asterisks = P<0.01.

### Mutants exhibited metabolic flux rewiring under acetate condition compared to WT

We performed a ^13^C tracer experiment involving *E.coli* WT and its mutant strains to determine the intracellular metabolic fluxes under acetate as a carbon source. A comprehensive network model (Supplementary materials 3) was defined wherein the labelling pattern of proteinogenic amino acids along with the physiological parameters (acetate uptake rate, NH3 uptake rate) were constrained to estimate the intracellular fluxes. These measured mass isotopomer distributions (MIDs) and the estimated metabolic fluxes within the accepted range of the SSRs (95% confidence) are provided in Supplementary materials 4. In Fig. 4, the normalized flux distribution maps (based on acetate uptake rate) for WT and its mutant strains were depicted. We identified the key nodes of acetate flux via central metabolic pathways in *E.coli,* as shown in Fig. 4A. The normalized acetyl-CoA flux splits towards the lipid synthesis (3%), TCA cycle pathway via citrate (67%), and glyoxylate shunt (30%). The net gluconeogenesis flux was estimated as 23% of the acetate uptake rate, of which 13% channelled through malic enzyme and 87% from OAA to PEP. Furthermore, about 3% of the acetate uptake rate routed through the PP pathway. At several nodes, such as α-KG, OAA, PEP, PYR, and 3PG, the carbon from acetate contributes towards amino acid biosynthesis, which amounted to ~33% of the acetate uptake rate.

**Fig. 4.**
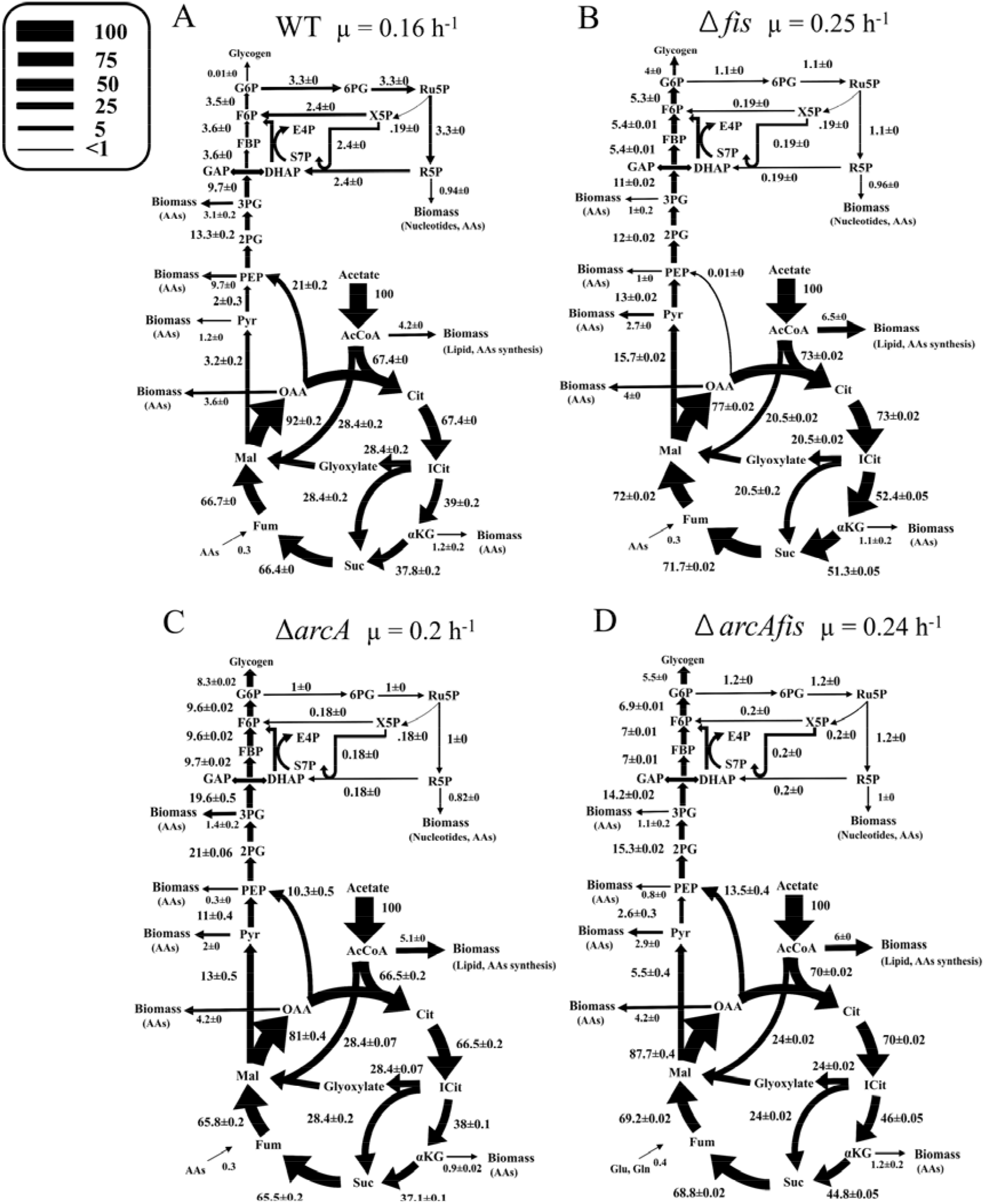
^13^C-Metabolic Flux distribution of acetate metabolism in *E.coli* MG1655 A) WT and mutants, B) Δ*fis*, C) Δ*arcA* and D) Δ*arcAfis*. All the intracellular fluxes are normalized with the 100 units of acetate uptake rate. Here, indicated errors reflect the standard deviation.

In the case of mutants, we observed the rewiring of fluxes in acetate metabolism. The normalized flux from acetyl-CoA towards citrate of 73% in Δ*fis* (Fig. 4B), 66.5% in Δ*arcA* (Fig. 4C) and 70% in Δ*arcAfis* (Fig. 4D) with no significant change in Δ*arcA*. Similarly, all mutants contributed higher flux towards fatty acid synthesis from 3% in WT to 5%, 4% and 4.5% in Δ*fis,* Δ*arcA* and Δ*arcAfis,* respectively. Fluxes from isocitrate to glyoxylate shunt decreased from 28% to 20% in Δ*fis* and 23% in Δ*arcAfis*. The net acetate flux towards amino acids also reduced to 20%, 22%, 21% in Δ*fis,* Δ*arcA* and Δ*arcAfis,* respectively (from 33% in WT). In Δ*fis*, the net gluconeogenesis flux decreased from 23% to 12.9% via malic enzyme with an insignificant amount from OAA to PEP. Instead, in Δ*arcA,* the net gluconeogenesis flux is 23.4%, of which 13.1% passed through malic enzyme and 10.3% from OAA to PEP. However, in Δ*arcAfis*, the net gluconeogenesis flux was 23%, similar to that seen in WT, but an alteration in the split was observed, i.e. about 10% flux passes through malic enzyme and 90% from OAA to PEP. In all the mutants, marginal flux ~1% of the acetate uptake rate was channelled through the PP pathway. Interestingly, mutants were characterized by carbon accumulation in the form of glycogen, with the highest in Δ*arcA* of ~ 8%, followed by Δ*arcAfis* and Δ*fis* of 5.5% and 4%, respectively.

In summary, we observed that mutants showed increased glycogen accumulation rate (4-8%), decreased flux towards amino acid biosynthesis (35 %) and decreased PP pathway fluxes (from 3% to 1 %).

### Δfis and ΔarcAfis liberated higher CO2 through the TCA cycle and anaplerotic pathway

Acetate metabolism in *E.coli* is majorly driven by high fluxes through the TCA cycle and gluconeogenesis, associated with CO_2_ efflux (Fig. 5A). A net CO_2_ release of 112%, 123%, 103% and 114% for WT, Δ*fis*, Δ*arcA* and Δ*arcAfis,* respectively, was observed. The significant proportion of CO_2_ generation was from TCA cycle via isocitrate to α-KG and α-KG to succinyl-CoA with fluxes of 77%, 104%, 75% and 90% for WT, Δ*fis*, Δ*arcA* and Δ*arcAfis*, respectively. In anaplerosis, malate to pyruvate and OAA to PEP are the significant pathways associated with CO_2_ efflux. Interestingly, CO_2_ flux from malate to pyruvate was remarkably higher in Δ*fis* (15%), followed by Δ*arcA* (13%) and Δ*arcAfis* (5.5%) compared to WT (3%). However, the CO_2_ fluxes decreased in OAA to PEP from 20% in WT to 10% in Δ*arcA* and 13.5% in Δ*arcAfis,* but no flux was reflected in Δ*fis.* Mainly, ~7% of the total CO_2_ efflux was evolved from the amino acid biosynthesis in WT and remarkably consistent of ~3% across all the mutants. The PP pathway involving the synthesis of Ru5P, generates CO_2_ but only in the ranges of 1-3 %. CO_2_ flux required for arginine biosynthesis was remarkably consistent ~ 0.3 % amongst all the strains.

**Fig. 5.**
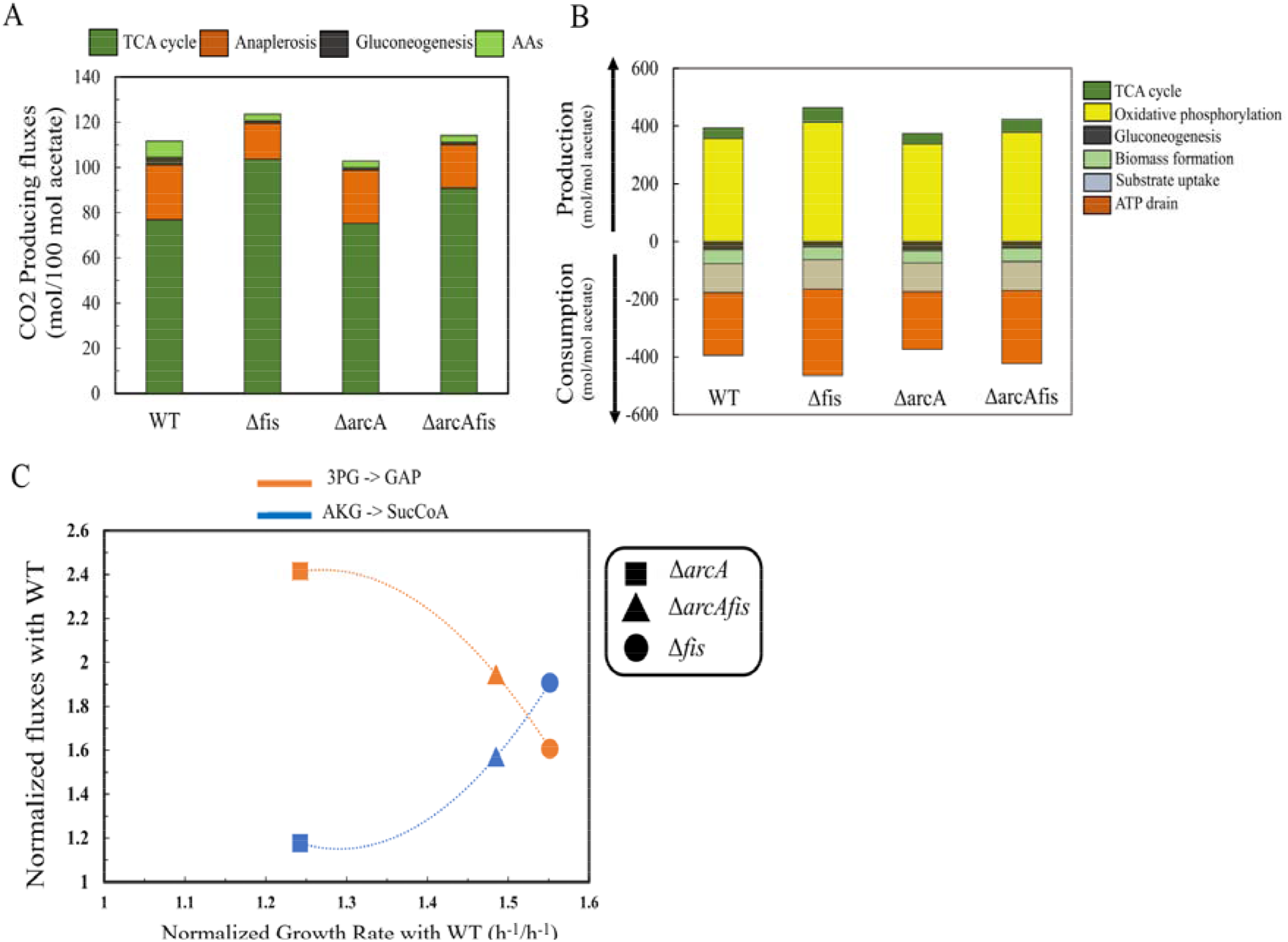
Key metabolic variations among the *E.coli* WT and mutants are shown under acetate condition. A) Respiratory CO_2_ efflux contributed by TCA cycle, anaplerosis, gluconeogenesis and amino acid biosynthesis B) ATP production and consumption estimated from ^13^C MFA and C) Variations of growth rate and fluxes representative of the TCA cycle reaction (AKG->SucCoA) in blue colour and Gluconeogenesis (3PG->GAP) fluxes in an orange colour described in mutants, normalized growth rate with WT, Δ*arcA* (1.25 h^−1^/ h^−1^), Δ*arcAfis* (1.48 h^−^ 1/ h^−1^) and Δ*fis* (1.55 h^−1^/ h^−1^).

Interestingly, the oxygen consumption rate was similar to the net release of CO_2_, indicating a respiratory coefficient of one in WT, which is in good agreement with the previously reported data (Andersent and Meyenburgt, 1980) and remained same in all the mutants. Hence, our findings demonstrate that the mutants (Δ*fis* and Δ*arcAfis*) exhibit an increased CO_2_ efflux through the TCA cycle and anaplerotic pathways in correlation with the increase in growth rate.

### Δfis and ΔarcAfis exhibit a higher trend of ATP production

Considering the normalized fluxes, the first committed step of acetate oxidation consumed 100 units of ATP flux across all the strains (Fig. 5B). The majority of ATP flux is utilized towards ATP maintenance which accounts for the difference in the total ATP synthesis (374-464%) and the consumption of ATP towards biomass (174-264%), with maximum costs in Δ*fis* and Δ*arcAfis* of 67% and 53% respectively compared to the WT with 34%. In OAA to PEP node, ATP utilization flux decreased by 50% in Δ*arcA* and 35% in Δ*arcAfis* than WT with negligible flux routes through this pathway in Δ*fis*. Fluxes through pyruvate to PEP enhanced by 6 folds in Δ*fis* and 5.5 folds in Δ*arcA* compared with WT. In the gluconeogenesis pathway (3PG to GAP), the ATP consumption rate was enhanced by 111% in Δ*arcA* and 46% in Δ*arcAfis* compared to the WT. In the TCA cycle, ATP production through succinyl-CoA synthetase accounts for 37%, 51%, 37% and 44% in WT, Δ*fis*, Δ*arcA* and Δ*arcAfis*, respectively. Oxidative phosphorylation contributed significantly towards ATP production, with 16% increase in Δ*fis*, 6% in Δ*arcAfis* and remained similar in Δ*arcA* compared to the WT. ATP utilization flux roughly remained the same towards biomass synthesis in all the strains. Hence, Δ*fis* and Δ*arcAfis* showed an enhancement in the total ATP production via the TCA cycle and oxidative phosphorylation but remained similar in Δ*arcA* than WT.

### Mutants exhibited high TCA cycle fluxes and variable gluconeogenesis and accumulation of glycogen

To compare the fluxes towards the TCA cycle and gluconeogenesis, the absolute flux values were determined from isocitrate to α-KG (extent of TCA) and 3PG to GAP (extent of gluconeogenesis) and plotted against the growth rate. Fig. 5C shows these values for the mutants relative to the absolute values seen in WT and plotted against the growth rate. It was observed that the flux towards the TCA cycle increases with an increase in growth rate and the net flux towards gluconeogenesis decreases with the growth rate. Our findings demonstrate that the TCA cycle flux is maximum in Δ*fis*, followed by the Δ*arcAfis* and Δ*arcA* strain, respectively. Note that the relative flux towards the TCA cycle in Δ*arcA* was like that in WT. This signifies that the higher absolute flux observed in Δ*arcA* strain (about 20%) was due to a 20% increase in acetate uptake rates. Whereas in the Δ*fis* strain, a 91% increase in the TCA flux relative to that in a WT is a combination of an increase in the acetate uptake rate (about 42%), and higher flux channelled towards the TCA cycle (49%). Δ*arcAfis* showed 57% higher flux in the TCA cycle than in WT, an intermediary to that seen in single mutants.

The higher flux towards the TCA cycle results in a higher ATP synthesis due to oxidative phosphorylation and directly correlates to higher growth rates (Fig. 5C). The extent of gluconeogenesis decreased with an increase in the growth rate. This suggested that in WT, the TCA cycle flux was limiting, and excess gluconeogenesis flux was achieved. Hence, the increased gluconeogenesis flux resulted in higher flux towards amino acids biosynthesis. The absolute flux towards gluconeogenesis was seen higher than in WT and was 140%, 94% and 60% higher in Δ*arcA*, Δ*arcAfis* and Δ*fis*, respectively. This implicit an overcompensation of flux towards gluconeogenesis in Δ*arcA*, which resulted in a higher amount of glycogen accumulation. Δ*fis* had a lower accumulation of glycogen, indicating a lower compensation, while in Δ*arcAfis*, the value is intermediatory of Δ*arcA* and Δ*fis*. In the case of WT, the absolute flux towards both the TCA cycle and flux towards gluconeogenesis was the lowest. Further, the PP pathway’s fluxes were lower in the mutants with 0.26, 0.2 and 0.25 (mmol/gDCW/h) in Δ*fis*, Δ*arcA* and Δ*arcAfis*, respectively compared to that in WT (0.53).

## Discussion

Global transcription factors play a significant role in controlling microbial metabolism under genetic and environmental perturbations. In this study, we deciphered metabolic phenotypes of *E.coli* regulatory mutants Δ*fis*, Δ*arcA* and double deletion mutant, Δ*arcAfis*, under acetate using steady-state ^13^C MFA. The global transcriptional factors *fis* and *arcA* are known to regulate acetate uptake rates by either directly regulating the uptake rate (*fis and arcA*) or by regulating the glyoxylate shunt and TCA cycle genes (*arcA*) (Browning et al., 2004; Perrenoud and Sauer, 2005; Seshasayee et al., 2012; Park et al., 2013; Iyer et al., 2021). Previous studies have reported the negligible effect of Δ*arcA* and Δ*fis* on the growth and metabolic phenotype of *E.coli* under glucose as a carbon source aerobically (Sagit Shalel Levanon et al., 2005; Shimizu et al., 2014). Mutants demonstrated higher acetate uptake rates along with the lower uptake rates of the nitrogen source. However, this phenomenon was anticipated in Δ*fis*, Δ*arcA* and Δ*arcAfis* as *fis* and *arcA* are known to regulate acetate uptake gene (*acs*) directly. Despite the fast growth exhibited by mutants, they displayed a lowered flux towards the amino acid biosynthesis (lowered by > 35%) and PP pathway relative to that observed in WT strain, indicating a lesser nitrogen utilization in mutants which indeed channelled more flux towards glycogen accumulation.

In the Δ*fis*, excess flux is channelled towards the TCA cycle in addition to the increased uptake rate of acetate, resulting in higher ATP synthesis. The lowered fluxes towards gluconeogenesis and ammonia uptake rate resulted in a better balance between energy and precursor synthesis, indeed resulting in a 56% higher growth rate but compromised with the carbon accumulation as glycogen. In the case of Δ*arcA* strain, an increased flux in the TCA cycle was mainly due to a higher acetate uptake rate, while lower ammonia uptake resulted in reduced amino acid flux. However, excess carbon resulted in increased glycogen accumulation. Previous studies have also reported that the cell induces glycogen storage to provide nutrients during stress condition (Wilson et al., 2010; Sekar et al., 2019). The increased growth rate was mainly due to an increased acetate uptake rate. The double deletion mutant, Δ*arcAfis*, demonstrated a combinatorial effect with the net state closer to Δ*fis* strain. It was interesting to note that the mutants demonstrated a better growth phenotype than WT. In the presence of glucose, it is well known that the acetate uptake is inhibited through the regulator *fis* and *arcA*. However, in a gluconeogenic carbon source such as acetate medium, it revealed a higher uptake rate and a better distribution of carbon and nitrogen towards energy and biomass synthesis, resulted in a better growth phenotype. Other global regulators, such as *crp, fnr* and *ihf,* are also known to perturb nitrogen uptake rate and its homeostasis (Pal et al., 2020). These effects are demonstrated in growth on a glucose medium. It was interesting to note that these effects had a beneficial result in the mutants while growing under acetate condition. Our study highlighted that the mutants had higher growth rates and acetate uptake rates with a lower ammonia uptake rate. ^13^C-MFA displayed lower gluconeogenesis and higher flux through the TCA cycle, which results in high energy status (ATP) in mutants. They exhibited a growth advantage to the system under acetate carbon sources by re-routing the fluxes where a high TCA cycle and low gluconeogenesis was preferred. Further, we observed that mutants channelled lower fluxes towards amino acids biosynthesis.

In summary, mutants exhibited metabolic readjustments to metabolize acetate via variations in the TCA cycle fluxes, glyoxylate shunt, anaplerosis, gluconeogenesis and glycogen accumulation resulting in higher growth and respiration. Carbon usage efficiency has enhanced in all the mutants with better fitness in growth and acetate utilization rate. This provide insights into the critical role of global transcriptional regulators *arcA* and *fis* that confer control over acetate metabolism in *E.coli.* The study demonstrates that in mutants, a better balance between energy and precursor synthesis through gluconeogenesis resulted in better growth, which lacked in the WT. Apart from understanding acetate metabolism, these findings also anticipated in rational metabolic engineering of strains to better utilize acetate towards targeted valuables for the biochemical synthesis of biofuels and various therapeutic products (Zhang et al., 2019).

## Author’s Contribution

S.J, K.V.V and S.K.M. conceived the idea, S.J and S.K.M designed the experiments; S.J and M.S.I designed the knockout strains; S.J. performed all the experiments. P.J assisted in the ^13^C GC-MS pipeline. S.J did the data mining. S.J, M.S.I, S.K.M, K.V.V analyzed the data. and K.V.V wrote the original draft. S.J, S.K.M and K.V.V reviewed and edited the paper.

## Acknowledgments

We acknowledge the BioX Centre and Central facility of IIT-Mandi for providing GC-MS and NMR facility. We thank Dr Sumana Srinivasan for her inputs in the manuscript. S.J. thank the Council of Scientific and Industrial Research (CSIR), Government of India (09/087(0857)/2016-EMR-I) for the fellowship.

