## Supplementary material for "Mutant strains of *Escherichia coli* lacking global regulators, *arcA* and *fis*, demonstrate better growth fitness by pathway reprogramming under acetate metabolism": Supp 1

**Supplementary Table 1: Primer design information used in the present study using one-step inactivation protocol.**

| **Name** | **Primer Sequence (5’-3’)** |
| --- | --- |
| **Primers used for generation of knockouts** | |
| ArcA KO fwd | CTTTTGTACTTCCTGTTTCGATTTAGTTGGCAATTTAGGTAGCAAAC GTGTAGGCTGGAGCTGCTTCG |
| ArcA KO rev | CGGCGCTAAAAAGCGCCGTTTTTTTTGACGGTGGTAAAGCCGA CCGGGGATCCGTCGACC |
| Fis KO fwd | GGCATACTTCGAAAATTTTGCGTAAACAGAAATAAAGAGCTGACAGAACT GTGTAGGCTGGAGCTGCTTCG |
| Fis KO rev | CCATGCCGAGTAGCGCCTTTTTAATCAAGCATTTAGCTAACCTGAA CCGGGGATCCGTCGACC |
| **External primers (200 bp upstream and downstream to the gene) used for detection of knockouts** | |
| ArcA Locus Fwd | TTTTGACACTGTCGGGTCCTGAGGGAAAGT |
| ArcA Locus Rev | TTGGGAACCAGTGTGCTGGTGGTGG |
| Fis Locus Fwd | GGAACTGCATGACTTTTATGGTCCGGCA |
| Fis Locus Rev | CAAAGATCAGATCGACACTTTCGGCGG |

**The E. coli variants used in the work are given as follows**

| **Strain Name** | **Genotype** | **Source** |
| --- | --- | --- |
| *E. coli* K12 MG1655 WT | F-, λ-, rph- | Keio collection *CGSC #6300* |
| *E. coli* K12 MG1655 *Δ*arca | F-, λ-, rph-, *Δarca kan* | This study |
| *E. coli* K12 MG1655 *Δ*fis | F-, λ-, rph-, *Δfis kan* | This study |
| *E. coli* K12 MG1655 *Δ*arca *Δ*fis | F-, λ-, rph-, *Δarca Δfis kan* | This study |
