## Supplementary material for "Mutant strains of *Escherichia coli* lacking global regulators, *arcA* and *fis*, demonstrate better growth fitness by pathway reprogramming under acetate metabolism": Supp 3

**Supplementary Information 3: Carbon transition network model employed in the INCA modeling for generating the ^13^C Flux maps in *E.coli* MG1655 and its mutant strains**

**Acetate catabolic Pathway**

1. Ac.ex (ab) -> Ac.in (ab)

2. Ac.in (ab) + ATP -> AcCoA (ab)

3. AcCoA (ab) ->AcCoA.ext (ab)

**TCA cycle**

| 4. OAA (abcd) + AcCoA (ef) -> Cit (dcbfea) |
| --- |
| 5. Cit (abcdef) -> ICit (abcdef) |
| 6. ICit (abcdef) -> GlyOxy (ab) + Suc (cdef) |
| 7. GlyOxy (ab) + AcCoA (ef) -> Mal (abef) |
| 8. ICit (abcdef) -> AKG (abcde) + CO2 (f) + NADPH |
| 9. AKG (abcde) -> SucCoA (bcde) + CO2 (a) + NADH |
| 10. SucCoA (abcd) ->Suc (abcd) + ATP |
| 11. Suc (abcd) -> Fum (abcd) + FADH2 |
| 12. Fum (abcd) -> Mal (abcd) |
| 13. Mal (abcd) -> OAA (abcd) + NADH |
| 14. Mal (abcd) -> Pyr (abc) + CO2 (d) + NADPH |

**Gluconeogenesis**

15. Pyr (abc) + ATP -> PEP (abc)

| 16. Pyr (abc) -> CO2 (a) + AcCoA (bc) + NADH |
| --- |
| 17. OAA (abcd) + ATP -> PEP (abc) + CO2 (d) |
| 18. PEP (abc) <-> 2PG (abc) |
| 19. 2PG (abc) <->3PG (abc) |
| 20. 3PG (abc) + ATP + NADH <-> GAP (abc) |
| 21. GAP (abc) <-> DHAP (abc) |
| 22. GAP (abc) + DHAP (def) -> FBP (abcdef) |
| 23. FBP (abcdef) -> F6P (abcdef) + ATP |
| 24. F6P (abcdef) -> G6P (abcdef) |
| 25. G6P (abcdef) ->Glycogen (abcdef) |
| **Pentose Phosphate Pathway** |
| 26. G6P (abcdef) -> 6PG (abcdef) + NADPH |
| 27. 6PG (abcdef) -> Ru5P (bcdef) + CO2 (a) + NADPH |
| 28. Ru5P (abcde) -> R5P (abcde) |
| 29. Ru5P (abcde) -> X5P (abcde) |
| 30. R5P (abcde) + X5P (fghij) <->S7P (fgabcde) + GAP (hij) |
| 31. S7P (nocdxyz) + GAP (pqr) <-> E4P (dxyz) + F6P (nocpqr) |
| 32. X5P (abcde) + E4P (fghi) <-> F6P (abfghi) + GAP (cde) |

| **Amino Acid Biosynthesis** | | |
| --- | --- | --- |
|  | 33. AKG (abcde) + NADPH + NH3 -> Glu (abcde) | |
| 34. | 34. Glu (abcde) + ATP + NH3 -> Gln (abcde) | |
|  | | 35. Glu (abcde) + ATP + 2 NADPH -> Pro (abcde) |
|  | | 36. Glu (abcde) + CO2 (f) + Gln (ghijk) + Asp (lmno) + AcCoA (pq) + 5 ATP + NADPH |
|  | | -> Arg (abcdef) + AKG (ghijk) + Fum (lmno) + Ac (pq) |
|  | | 37. OAC (abcd) + Glu (efghi) ->Asp (abcd) + AKG (efghi) |
|  | | 38. Asp (abcd) + 2 ATP + NH3 -> Asn (abcd) |
|  | | 39. Pyr (abc) + Glu (defgh) ->Ala (abc) + AKG (defgh) |
|  | | 40. Ser (abc) -> Pyr (abc) + NH3 |
|  | | 41. 3PG (abc) + Glu (defgh) -> Ser (abc) + AKG (defgh) + NADH |
|  | | 42. Ser (abc) <-> Gly (ab) + MEETHF (c) |
|  | | 43. Gly (ab) <-> CO2 (a) + MEETHF (b) + NADH + NH3 |
|  | | 44. Thr (abcd) <-> Gly (ab) + AcCoA (cd) + NADH |
|  | | 45. Ser (abc) + AcCoA (de) + 3 ATP + 4 NADPH + SO4 -> Cys (abc) + Ac (de) |
|  | | 46. Asp (abcd) + Pyr (efg) + Glu (hijkl) + SucCoA (mnop) + ATP + 2 NADPH ->  LL-DAP (½ abcdgfe + ½ efgdcba) + AKG (hijkl) + Suc (½ mnop + ½ ponm) |
|  | | 47. LL-DAP (½ abcdefg + ½ gfedcba) -> Lys (abcdef) + CO2 (g) |
|  | | 48. Asp (abcd) + 2 ATP + 2 NADPH -> Thr (abcd) |
|  | | 49. Asp (abcd) + METHF (e) + Cys (fgh) + SucCoA (ijkl) + ATP + 2 NADPH ->  Met (abcde) + Pyr (fgh) + Suc (½ ijkl + ½ lkji) + NH3 |
|  | | 50. Pyr (abc) + Pyr (def) + Glu (ghijk) + NADPH -> Val (abcef) + CO2 (d) + AKG  (ghijk) |
|  | | 51. AcCoA (ab) + Pyr (cde) + Pyr (fgh) + Glu (ijklm) + NADPH ->  Leu (abdghe) + CO2 (c) + CO2 (f) + AKG (ijklm) + NADH |
|  | | 52. Thr (abcd) + Pyr (efg) + Glu (hijkl) + NADPH -> Ile (abfcdg) + CO2 (e) + AKG  (hijkl) + NH3 |
|  | | 53. PEP (abc) + PEP (def) + E4P (ghij) + Glu (klmno) + ATP + NADPH -> Phe  (abcefghij) + CO2 (d) + AKG (klmno) |
|  | | 54. PEP (abc) + PEP (def) + E4P (ghij) + Glu (klmno) + ATP + NADPH ->  Tyr (abcefghij) + CO2 (d) + AKG (klmno) + NADH |
|  | | 55. Ser (abc) + R5P (defgh) + PEP (ijk) + E4P (lmno) + PEP (pqr) + Gln (stuvw) + 3  ATP + NADPH -> Trp (abcedklmnoj) + CO2 (i) + GAP (fgh) + Pyr (pqr) +  Glu (stuvw) |
|  | | 56. R5P (abcde) + FTHF (f) + Gln (ghijk) + Asp (lmno) + 5 ATP ->  His (edcbaf) + AKG (ghijk) + Fum (lmno) + 2 NADH |
| **One-Carbon Metabolism** | | |
|  | | 57. MEETHF (a) + NADH -> METHF (a) |
|  | | 58. MEETHF (a) -> FTHF (a) + NADPH |

**Oxidative Phosphorylation**

59. NADH + ½ O2 -> 2 ATP

60. FADH2 + ½ O2 -> 1 ATP

| **ATP Hydrolysis** | |
| --- | --- |
|  | 61. ATP -> ATP:ext |
| **Transport** | |
|  | 62. CO2 (a) -> CO2.ext (a)  63. CO2.unlabeled + CO2 -> CO2 + CO2.out |
|  | 64. O2.ext -> O2 |
|  | 65. NH3.ext -> NH3 |
|  | 66. SO4.ext -> SO4 |
| **Transhydrogenase Reaction**  67. NADH <-> NADPH  **Biomass Formation** | |
|  | 68. 0.488 Ala + 0.281 Arg + 0.229 Asn + 0.229 Asp + 0.087 Cys + 0.250 Glu + 0.250 Gln + 0.582 Gly + 0.090 His + 0.276 Ile + 0.428 Leu + 0.326 Lys + 0.146 Met + 0.176 Phe + 0.210 Pro + 0.205 Ser + 0.241 Thr + 0.054 Trp + 0.131 Tyr + 0.402 Val + 0.205 G6P + 0.071 F6P + 0.754 R5P + 0.129 GAP + 0.619 3PG + 0.051 PEP + 0.083 Pyr + 2.510 AcCoA + 0.087 AKG + 0.340 OAC + 0.443 MEETHF + 33.247 ATP + 5.363 NADPH -> 39.68 Biomass + 1.455 NADH |
